# Tuning Intermediate Filament Mechanics by Variation of pH and Ion Charges

**DOI:** 10.1101/784025

**Authors:** Anna V. Schepers, Charlotta Lorenz, Sarah Köster

## Abstract

The cytoskeleton is formed by three types of filamentous proteins – microtubules, actin filaments, and intermediate filaments (IFs) – and enables cells to withstand external and internal forces. Vimentin is the most abundant IF protein in humans and assembles into 10 nm diameter filaments with remarkable mechanical properties, such as high extensibility and stability. It is, however, unclear to which extent these properties are influenced by the electrostatic environment. Here, we study the mechanical properties of single vimentin filaments by employing optical trapping combined with microfluidics. Force-strain curves, recorded at varying ion concentrations and pH values, reveal that the mechanical properties of single vimentin IFs are influenced by pH and ion concentration. By combination with Monte Carlo simulations, we relate these altered mechanics to electrostatic interactions of subunits within the filaments. We thus suggest possible mechanisms that allow cells to locally tune their stiffness without remodeling the entire cytoskeleton.

## 1 Introduction

Biological cells are constantly exposed to varying intra- and extra-cellular forces, thus their mechanical properties are challenged. It is meanwhile well accepted that the mechanical properties of cells are, to a large extent, governed by the cytoskeleton, a stabilizing, yet flexible framework, which is composed of microtubules, actin filaments, and intermediate filaments (IFs), together with motor proteins and crosslinkers. While microtubules and actin filaments are conserved in eukaryotic cell types, there are around 70 different genes encoding IFs in man that are expressed according to the cell’s specific requirements. ^1,2^ Despite differences in their amino acid sequence, all cytoskeletal IF proteins share their secondary structure with a tripartite alpha-helical rod and disordered head and tail domains. ^3,4^ During the hierarchical assembly, two monomers form a parallel coiled coil and these dimers organize into anti-parallel tetramers in a half-staggered arrangement. The tetramers rapidly assemble laterally to form unit length filaments (ULFs) within 100 ms, ^5^ which in turn elongate to filaments by end-to-end annealing, ^3^ The resulting biopolymer comprises a complex high-order arrangement of coiled coils, ^3^ which allows IFs to be extended up to at least 4.5-fold their initial length, ^6,7^ including a strong loading-rate dependence. ^8^ This enormous extensibility is in stark contrast to microtubules and F-actin. ^9^

Vimentin is the IF typically expressed in mesenchymal cells ^2^ and up-regulated during the epithelial-to-mesenchymal transition in wound healing, early embryogenesis, and cancer metastasis. ^10^ In addition to being highly extensible ^8,11,12^ vimentin is flexible ^13–15^ and stable. ^16^ While the extensibility describes how much a filament can be elongated before rupture, it does not explain the mechanisms underlying the extension and therefore the response of the filament to an applied strain, *i.e.* the relative elongation of the filament. By analyzing force-strain curves, fundamental properties of the material are determined. For example, the elasticity and the Young’s modulus can be derived from a linear force increase. During stretching, three regimes are observed in the force-strain data: ^8,11,12,17,18^ an initial linear, elastic increase, a plateau of relatively constant force, and a subsequent stiffening regime. These regimes have been linked to structural changes in the protein. ^17^ The initial linear increase is described as elastic stretching of the filament consisting of the alpha helices, the plateau as the unfolding of alpha-helical structures, and the stiffening as the further stretching of the unfolded structure. Once the filament is stretched beyond the linear regime, the deformation becomes inelastic. The three-regime behavior is, among the cytoskeletal filaments, unique for IFs, as F-actin and microtubles rupture at much lower strains. ^9^ The IFs show a pronounced softening upon re-stretching, while, at the same time, the elongation itself is fully reversible and the noise threshold of 2 pN is reached at residual strains of 0.05*±*0.02 (mean, standard deviation). ^12,19^

A similar three-regime force-distance behavior has been found for single coiled coils, ^20,21^ which are a common theme in protein structures. The overall structural stability of coiled coils depends on the buffer conditions, however, the results are partially conflicting. Some studies show an increased stability at low pH compared to neutral pH, ^22,23^ whereas others find higher stability in neutral pH conditions. ^24,25^ It has been discussed that the response of a coiled coil to altered pH conditions depends on the concentration of salt ions in the buffer but also on the sequence of the peptide. ^26^ Molecular dynamics simulations of the isolated vimentin coiled-coil dimer ^18^ agree qualitatively with experimental force-strain curves of single coiled coils ^20,21,27^ and vimentin filaments. ^8,12^ These results indicate that coiled coils play a pivotal role in the force-strain response of mature IFs.

Here we address the open question of how strongly the variability of the mechanical response observed for coiled coils in different measurement conditions is conserved in fully assembled vimentin IFs. We study the response of mature vimentin IFs to tuning of the ionic conditions of the buffer and to the internal charge distribution in the protein by adjusting the pH of the buffer, and find that both factors strongly influence the mechanics of single vimentin IFs. Monte-Carlo simulations enable us to link electrostatic interactions within the filament and modifications of the free energy landscape of the unfolding reaction to the observed force-strain behavior. These results extend the list of remarkable mechanical properties of IFs by adding a responsiveness to pH and ionic environment to the established loading rate sensitivity ^8^ and tensile memory. ^12^ The possibility to tune filament mechanics fast and locally may have important implications within living cells.

## 2 Experimental

### 2.1 Experimental Design

The experiments were performed using a setup that combines optical traps, confocal microscopy and microfluidics (C-Trap, Lumicks, Netherlands). A four-inlet glass microfluidic flow cell enabled easy change of buffer during the experiment as the subchannels in the flow cell were separated by laminar flow. The syringes feeding the microfluidic flow cell were driven by air pressure. Solutions were injected into the four channels (the channel geometry is shown in Fig. S1a†) as follows: 1: beads in measuring buffer, 2: measuring buffer, 3: assembly buffer, 4: vimentin in assembly buffer. Figure 1b shows a simplified s ketch, excluding channel 2 that is used for calibration of the traps. For the optical trap measurements, the filaments were diluted 150-fold in assembly buffer. Maleimide-functionalized polystyrene beads (Kisker Biotech, Steinfurt, Germany) ^12,28^ were diluted in measuring buffer. The optical trap was calibrated by analysis of the power spectral density of the thermal fluctuations of the trapped beads. Filaments were tethered to trapped beads in assembly buffer in flow. The flow was then stopped, the beads with the tethered filament were moved to the measuring buffer and the filaments incubated for 30 seconds without flow. The filaments were stretched at a loading rate of 0.21 *±* 0.05 *μ*m/s. Filaments were stretched until rupture or until the force exceeded the trap potential and one of the beads was pulled out of the trap. For each condition, force-strain curves of at least seven single, stable filaments were recorded. Depending on the initial length of the filament, one measurement was completed within 30-60 s. As measuring buffers we used 2 mM PB at varying pH (5.8 - 8.5) and concentrations of KCl (0, 50, 100, 150 mM) or MgCl_2_ (0, 5, 10 mM) (Carl Roth, Germany). The physiological KCl concentration up to 150 mM, ^29^ which corresponds to the upper *c*(KCl) used here. The highest *c*(MgCl_2_)=10 mM used in this study is the threshold concentration observed for filament bundling and collapse of vimentin networks. ^30–32^ Albeit the pH in the extracellular space can be lower than 5.8, all lower values would be outside the buffer range of PB.

**Fig. 1.**
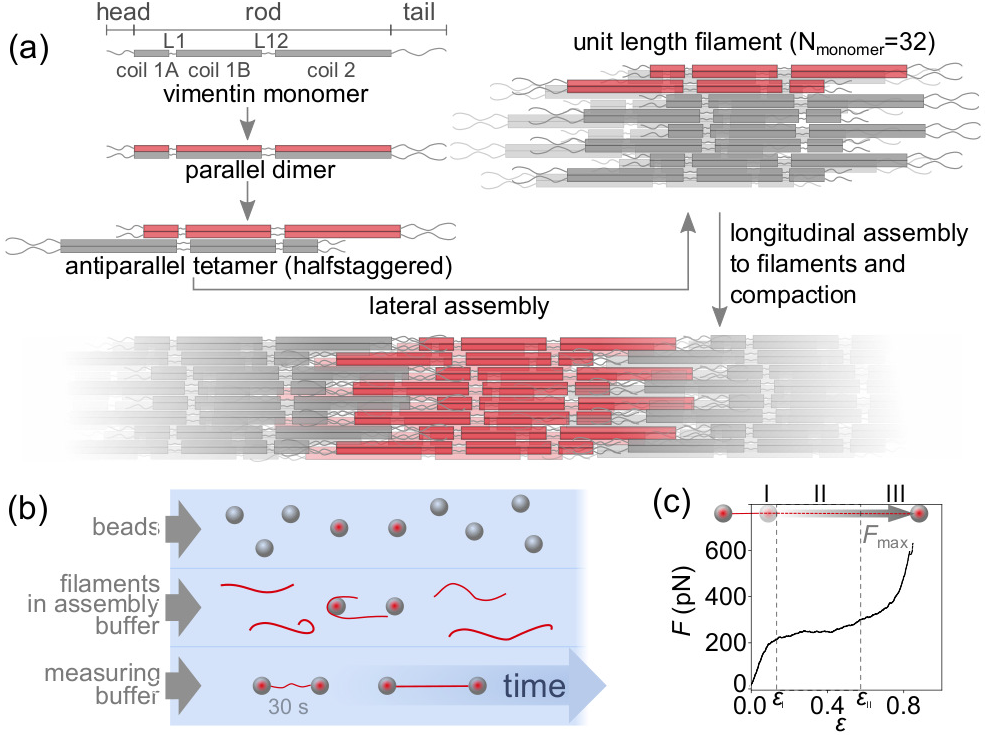
Stretching experiments on fully assembled vimentin filaments. (a) Schematic of the hierarchical assembly of vimentin monomers into filaments. In each step, the respective precursor subunit is highlighted in red. (b) Simplified measurement protocol of the optical trap experiment in microfluidic flow channels: Two beads are captured and calibrated (1), a single filament is covalently attached to both beads (2) and stretched after a 30 s incubation period in the measuring buffer (3). (c) Typical force-strain curve for a vimentin filament showing the elastic (I), plateau (II) and stiffening (III) regime. The strain at the transition of region I to II is *ε*_I_, from II to III it is *ε*_II_.

#### 2.1.1 Protein preparation

Human vimentin (mutant C328N with three additional amino acids, GGC, at the C-terminus) was recombinantly expressed and purified as previously described. ^12^ The protein was labeled with ATTO647N maleimide (ATTO-Tec, Germany), ^33,34^ Reconstitution and assembly was performed as previously reported. ^12^ In brief, unlabeled and labeled monomers were mixed (4% labeling ratio) and reconstituted at room temperature by dialysis into 2 mM phosphate buffer (PB, Carl Roth, Germany) at pH 7.5, decreasing the Urea (Carl Roth, Germany) concentration every 30 min in steps of 6, 4, 2, 1, 0 M Urea and an subsequent overnight dialysis step at 8-10°C. Filaments were assembled by dialysis into assembly buffer (2 mM PB with 100 mM KCl (Carl Roth, Germany) at pH 7.5) at 36°C for 16 h at a protein concentration of 0.2 g/L.

### 2.2 Data analysis

#### 2.2.1 Calculation of single force-strain curves

For each filament, the initial length, *L*_0_, was determined and used for the calculation of the strain: *ε* = (*L−L*_0_)*/L*_0_. To obtain *L*_0_, the raw force-distance curve was smoothed with a moving average with a window width of 10 data points to account for fluctuations of the trap. The initial length was set as the length at the last data point before the smoothed curve reached 5 pN.

#### 2.2.2 Analysis of the filament stability

Force-strain curves were plotted and the curves that showed a plateau and subsequent stiffening were categorized as stable filaments. All other filaments are categorized as instable. From confocal videos of the filament stretching process and the force-strain behavior of the stable filaments, bundles were identified and the force-strain curves of the bundles excluded from further analysis. Examples for confocal images of bundles and single filaments are shown in Fig. S2†.

#### 2.2.3 Calculation of average force-strain curves

For each condition, the average maximum strain of all stable filaments was calculated. Each stable force-strain curve was scaled to the average maximum strain, interpolated to 200 values, and the forces were averaged.

#### 2.2.4 Analysis of the slope of the plateaus

The plateau of each single force-strain curve was analyzed for all stable filaments. A typical analysis for one single force-strain curve is shown in Fig. S3†. The point of maximum strain, *ε*_max_, of each curve was used to find the mid data point, *ε*_mid_, of the curve (yellow in Fig. S3a†). The data points with relatively constant slope were determined. To do so, the differential of each single force-strain curve was calculated. To account for changes in length of the curve and noise of the data, each differential force-strain curve was then smoothed using a moving average with the width of 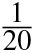 of the number of data points in the curve before *ε*_max_. The values 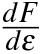 at the 10 data points before and after *ε*_mid_ were averaged to find 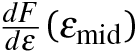. Next, the first maximum of 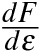 position A, was calculated (marked in F(ig. S3a †). The first and last data point to fulfill 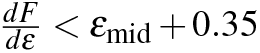.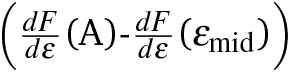 were termed *ε*_I_ (red, solid) and *ε*_II_ (blue, solid), respectively. To ensure that the transition regions between I, II and III were not included in the analysis of the slope of the plateau, a linear regression (green line) of the force-strain curve was calculated for the center 80% of the length of the plateau (between the open red and blue circles). The mean slope and standard deviation were calculated for each measuring condition.

#### 2.2.5 Analysis of the force of the plateaus

For the analysis of *F*_plateau_, *ε*_I_ determined from the average force-strain curve for each condition was used. The force at *ε*_I_ for each single force-strain curve was determined and the mean value and standard deviation were calculated.

#### 2.2.6 Determination of the end points of the elastic and plateau regions

For the determination of the slope of the plateau, the regression was calculated over a large region of the force-strain curve. This approach compensated for noise in the data. However, the noise limits the accurate determination of *ε*_I_ and *ε*_II_. Therefore, for the analysis of *ε*_I_ and *ε*_II_ for each condition, the average force-strain curves were used as shown in Fig. 2. The average curves were treated in the same way described above for the single force-strain curves. The only difference was that here the 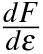 curves were smoothed with a moving average of 15 data points.

**Fig. 2.**
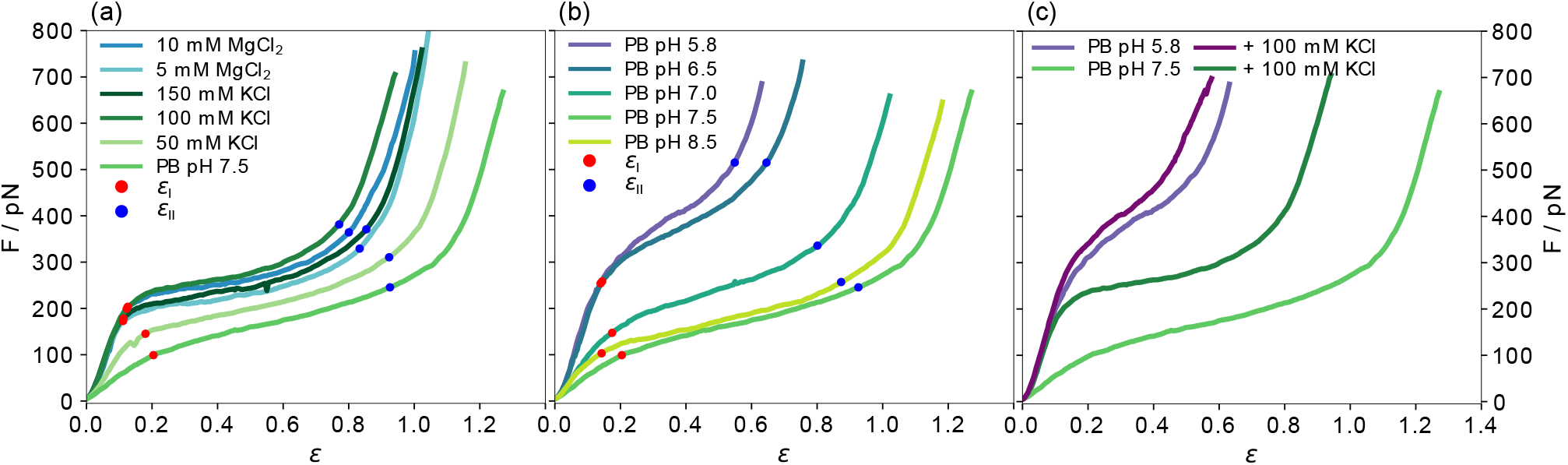
Force-strain behavior of single vimentin filaments. All curves shown are averages of the individual measurements shown in Figs. S4 and S5. The strain values at the end of the initial linear regime *ε*_I_ (red) and the plateau regime *ε*_II_ (blue) for all average curves are indicated. (a) Effect of indirect charge shifts caused by salt ions in the measurement buffer. (b) Effect of direct charge shifts by varying pH conditions. While *ε*_I_ is similar in all measurements, *ε*_II_ increases for lower *c*(KCl) and for increasing pH. (c) Comparison of the effect of an addition of 100 mM K^+^ ions at pH 7.5 and 5.8.

#### 2.2.7 Analysis of the initial slope of the force-strain curves

The initial slope was determined for strains from 0.02 to 0.1 or to a force of 100 pN, whichever occurred earlier.

### 2.3 Force-strain Monte-Carlo simulations

A vimentin filament was mechanically modeled as previously described. ^12,35^ The force-strain behavior of the modeled filament was determined by a Monte-Carlo simulation written in MatLab. ^35^ The simulation was run with one varied parameter while keeping the others constant as shown in Fig. 4a,b,e,f. The default parameters were the alpha-helical spring constant (*κ_α_* = 5.5), the number of monomers in a subunit (*N_sub_* = 32), the free energy difference between the alpha state and unfolded state (∆*G* = 2) and the length by which an alpha helix can extend upon unfolding ∆*L* = 1.

## 3 Results and discussion

### 3.1 Cations stiffen single vimentin IFs

To investigate the effect of the ionic environment on the mechanical behavior of single vimentin filaments, we stretch the filaments after incubation in different buffers. For comparability, all filaments are assembled in standard assembly buffer (2 mM phosphate buffer (PB) with 100 mM KCl, pH 7.5, see Fig. 1a for a schematic representation of the hierarchical assembly pathway) before the stretching experiment. A single filament is then tethered to optically trapped beads and incubated in the respective measuring buffer for 30 s before stretching. A graphical protocol of the experiment is shown in Fig. 1b. Fig. 1c shows a typical resulting force-strain curve, where the strain is defined as *ε = (L−L_0_)/L_0_* with the original filament length *L*_0_ measured 5 pN force.

The mechanical response of single filaments to stretching shows a clear dependence on the experimental conditions, as shown in Fig. 2. All panels show average data and the individual curves are omitted here for clarity, but are shown in Figs. S4 and S5. The three regimes that have been previously reported ^8,11,12,17,18^ are evident in the force-strain data recorded under standard assembly conditions as shown in Fig. 2a (100 mM KCl, see legend for color code): The initial linear increase (I in Fig. 1c), the plateau (II) and the subsequent stiffening at high strains (III) can be clearly distinguished. Note that what we describe as “plateau” here does not necessarily have a slope of zero, but a considerably decreased slope compared to the rest of the curve. The curves recorded at high salt concentrations, *i.e. c*(KCl) = 100 mM or 150 mM and *c*(MgCl_2_) = 5 mM or 10 mM, are consistent with this mechanical behavior. In particular, the initial slopes, which describe the initial elasticity of the filament, agree well between these four salt conditions, as shown in detail by the solid circles in Fig. 3a. These values are determined by fitting the initial slopes of the individual force-strain curves (Figs. S4 and S5†).

**Fig. 3.**
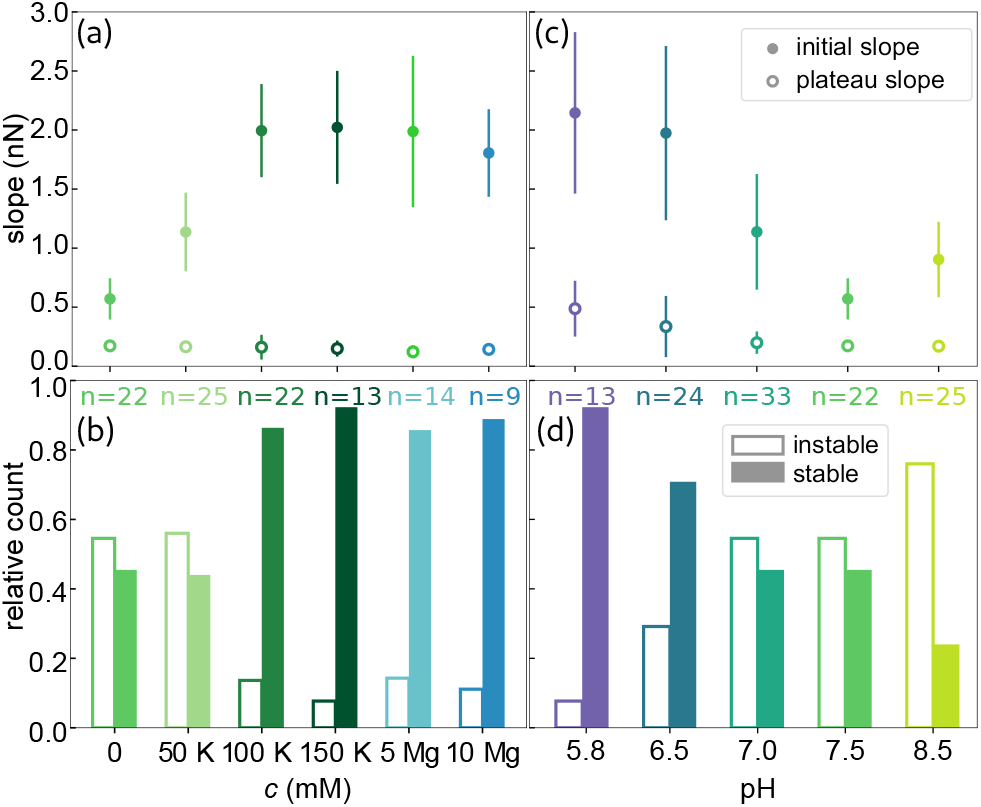
Mechanical properties of single vimentin filaments. (a),(c) Analysis of the initial slopes (solid circles) and slopes of the plateau regions (open circles) of all stable filaments. The error bars indicate the standard deviation. (b),(d) The stability of the measured filaments is presented as fractions of instable and stable filaments.

**Fig. 4.**
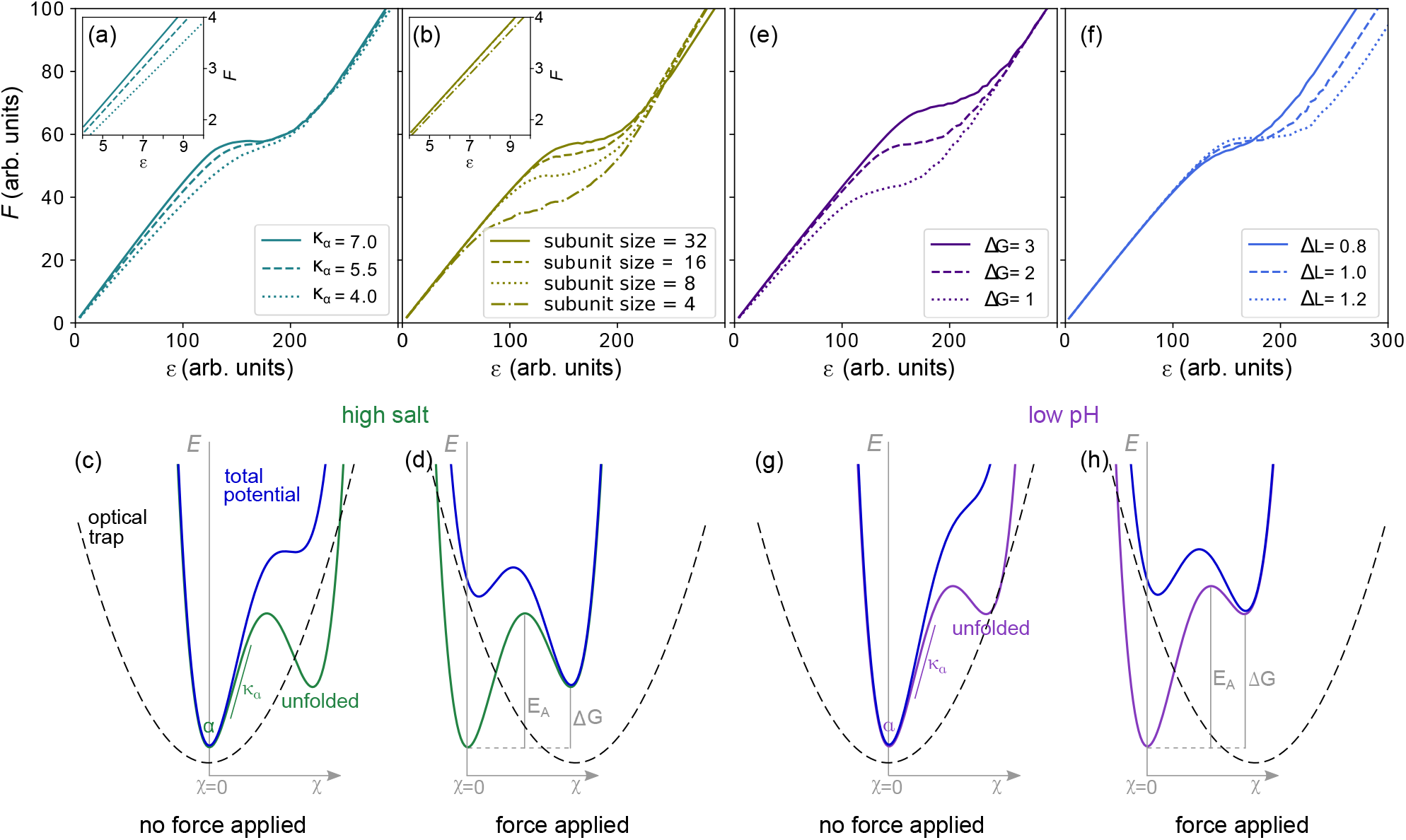
Monte-Carlo simulations of force-strain curves and schematics of energy landscapes. (a) An increased *κ_α_* causes an increase of the initial slope. (b) A stronger coupling into larger subunits moves the plateau to higher forces, decreases the slope of the plateau and weakly influences the initial slope. The insets in a and b show the initial slope for each parameter set. (c) Energy landscape *E* plotted against the reaction coordinate *χ* with minima for the alpha and unfolded state at high salt conditions (green) within the harmonic potential corresponding to the optical trap (dashed line). The resulting total potential is shown in blue. Without applied load, the alpha state is stable. (d) By moving the optical trap, and thereby the harmonic potential, the energy barrier is reduced and the unfolded state becomes more probable. (e) A higher free energy difference Δ*G* between the alpha and unfolded state increases *F*_plateau_ without affecting the slope of the plateau. (f) Increasing the length of the unfolded monomer increases *ε*_II_. **(G)** The suggested energy landscape at low pH, which leads to an increased energy barrier Δ*G*, shows a higher *E_A_* making the transition to the unfolded state less probable, (h) even after applying the same trap load as in (f).

When the filaments are incubated in low salt buffer (PB, pH 7.5), where tetramers are known to be stable, ^36^ before stretching, the mechanics change significantly. Fig. 2a shows that the complete curve is shifted to lower forces, the initial slope is lower and the plateau is less pronounced. The decreased initial slope is indicative of a softer material. As a consequence of this softening, the filament can be stretched to higher strains as compared to high salt buffers before the maximum force of the optical trap is reached. The curve measured at 50 mM KCl lies between the data for low salt buffer and the standard assembly buffer curve, as does the maximum strain for this condition. Independent of the measuring conditions, the strain at which the initial linear increase ends is at *ε*_I_ = 0.15 *±* 0.04 (see Fig. 2a), showing that the elastic extensibilty of the filament is not affected by the salt ions. It is remarkable that the slope of the plateau is constant for all buffers, as shown in Fig. 3a by the open circles.

Amino acids, the building blocks of proteins, may be either hydrophobic, polar or charged. Ions in the buffer interact with those polar and charged amino acids that are accessible within a supramolecular structure, thus mediating interactions within the assembled filament. Such electrostatic interactions caused by ions in the buffer can be regarded as *indirect* charge effects. The similarity between the curves at *c*(KCl) = 100 mM or 150 mM and *c*(MgCl_2_) = 5 mM or 10 mM indicates that predominantly the cations are causing the different behavior and not the Cl^−^ anions. In contrast, if it were mainly the Cl^−^ ions that caused the stiffening of the filaments, we would expect the curve at *c*(KCl) = 50 mM to lie above both MgCl_2_ curves since 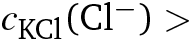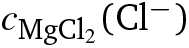 for all measured buffers.

The cations do not only stiffen but also stabilize the single filaments as shown in Fig. 3b by the relative count of stable filaments (solid bars) and instable filaments (open bars). The stability is determined from the shape of the force-strain curve. Filaments that break before the stiffening regime (III in Fig.1c) is reached, are classified as instable. The fraction of stable filaments increases from 0.45 for the measurements in low salt buffer to 0.88 *±* 0.02 (average and standard deviation from data at *c*(KCl) = 100 mM and 150 mM, and *c*(MgCl_2_) = 5 mM and 10 mM) at higher salt concentrations. The cations additionally promote bundling (see Fig. S2a†and Fig. S6a†). In particular, at *c*(MgCl_2_) = 10 mM, the amount of bundling limits the number of measurable single filaments. ^30,32^ This observation agrees well with the reported behavior of vimentin networks where – with increasing concentrations of multivalent ions – freely fluctuating networks collapse to dense aggregates. ^31^ It has further been reported that reconstituted vimentin networks stiffen upon addition of divalent ions, ^14,37,38^ and that the network stiffness increases with increasing concentration of divalent ions until total collapse of the network. ^39^ Combined with these findings from literature, our results show that the divalent ion dependent stiffening of vimentin networks is solely due to stronger interactions between filaments and not to additional stiffening of single filaments in the network. The small number of instable filament we observe at physiologically relevant *c*(KCl) ^29^ as well as all measured *c*(MgCl_2_) probably do not contribute considerably to cell mechanics. In line with the known high stability under large applied strains of IF networks, ^9^ and cell experiments that identify IFs as the load bearing elements at high strains, ^40^ our data thus suggest that the stable filaments determine IF mechanics in cells.

### 3.2 IF mechanics adapt to pH changes

Whereas the interactions of ions with the protein described in the previous section represent an indirect charge effect on the filament, we can also *directly* manipulate the charge of specific amino acids, *e.g.* by varying the pH of the buffer. To keep the two effects separate, at first, we use the curve recorded in the low salt buffer (2 mM PB, pH 7.5) as a starting point and do not add any additional salt ions. As the cytoplasmic pH in eukaryotic cells is reported to lie between 7.0 and 7.5, ^41,42^ we lower the pH to 7.0. The resulting curves are shifted to higher forces and the plateau region is shorter compared to the low salt buffer at pH 7.5 (see Fig. 2b and Fig. S5c,d†). The stiffening effect is amplified at even lower pH as shown by data for pH 6.5 and pH 5.8. The curves recorded in these low pH conditions are remarkably similar to each other and the filaments are considerably stiffer than in all previous measurement conditions with the plateau almost disappearing. If we, in contrast, increase the pH from 7.5 to 8.5, the mechanics of the filament do not change. Thus, the adaption of the filament occurs between pH 6.5 to 7.5 and it assumes an intermediate state at pH 7.0.

In line with the observations for varying salt concentrations reported above, the strain reached with the initial, linear increase (*ε*_I_= 0.16*±*0.03) is not strongly influenced by the pH of the measurement buffer. This result shows that the elastic extensibility of the filament is neither strongly affected by pH or ionic charge variation. Additionally, the initial slope at low pH is the same as for the standard assembly buffer or high salt concentrations (Fig. 3a,c), indicating that low pH and high salt have a similar effect on the initial stiffness of the filament. The decrease of the initial slope with increasing pH is plotted by solid circles in Fig. 3c and is evident in the force-strain curves at low strains in Fig. 2b. In contrast to the cations, the pH clearly affects the unfolding mechanism responsible for the plateau formation and the increased slope of the plateau region at low pH suggests that the force needed for unfolding events increases with decreasing pH. Because the onset of the stiffening moves to lower strains for low pH, the strain range of the plateau becomes very short. Unlike for filaments in varying salt conditions (Fig. S7a†), the ratio of the initial slope and slope of the plateau does not change with the pH (Fig. S7b†).

Taken together, our force-strain data on vimentin IFs at different pH values show that the overall stiffness of the filament and the unfolding are altered considerably when the charge of a few specific amino acids is varied. This assumption is further supported by the stability behavior of the filaments (see Fig. 3d). Here, the fraction of stable filaments, represented by solid bars, decreases with increasing pH, from 0.92 at pH 5.8 to 0.24 at pH 8.5, while the force-strain curves do not change between pH 7.5 and 8.5 or between 5.8 and 6.5. A stabilization or destabilization therefore still continues even if the force-strain behavior of the stable filaments are not affected. While the measuring condition clearly affects the stability of the filaments, there is no correlation of the stability, *i.e.* the maximum force reached during the experiment, and the initial length of the filament, as shown in Fig. S8†.

### 3.3 IF stiffening saturates at low pH

We observe an overall weaker stiffening by cationic (indirect) charge shifts compared to pH (direct) charge shifts on single vimentin filaments. This phenomenon is especially prominent in the plateau regions. To understand the interplay of the two effects, we compare two sets of data recorded at different pH (7.5 and 5.8) without additional salt and with 100 mM KCl each. The four average curves are shown in Fig. 2c. At pH 5.8, 100 mM KCl does not have a strong effect, and the curves with and without additional salt are strikingly similar (purple), especially when compared to the pronounced effect of 100 mM KCl at pH 7.5 (green). This observation suggests that the maximum stiffness has already been reached at low pH without salt and the ions only have a negligible effect. To ensure that the saturation of the stiffening effect we observe at low pH is constant on the time scales accessible here, we further investigate the temporal evolution of the adaptation of the mechanics of the filament at low pH. The negligible difference between curves (Fig. S9†) recorded at incubation times of 15 s, 30 s, and 60 s shows that the additional interactions within the filaments have developed already after a few seconds.

### 3.4 Variations in the free energy landscapes influence filament mechanics

The observed filament softening in low salt buffer and stiffening at low pH raises the question of how these mechanical properties are governed by molecular charge interactions within the filament. To answer this question, we first regard the initial slope of the force-strain curves in Fig. 2. The initial slope decreases when fewer monovalent cations are present and increases with decreasing pH.

We model force-strain curves by Monte Carlo simulations that are based on the hierarchical structure of the filaments. ^8,35^ In the model, monomers consist of a spring and an extendable element that can either be open or closed. The spring accounts for the elastic contribution of the alpha helices. Lateral arrangement of 32 such monomers represents one ULF. The filament is simulated as a series of 100 ULFs that are connected by springs. In the model, we can define the elastic properties of the springs *κ_α_*, the strength of the coupling between lateral monomers in each ULF, the free energy difference of the closed, *α*, and unfolded state Δ*G* and the length that is added by stretching a monomer Δ*L*.

In our 1D experimental setting, we can interpret the initial slope as a measure of the filament stiffness, which for the sake of modeling we describe by the spring constant of the filament, *κ_f_*. We expect an increase with the number of monomers, *N*, per cross-section of the filament. ^43^ We can, however, exclude the possibility of a reorganization of mature filaments *in vitro* by addition or loss of subunits as it only occurs on timescales of tens of minutes ^34^ which is much slower than the time scales of our experiments. We can therefore safely assume that the number of monomers is constant during our experiments. Instead, stiffening of the filament may originate from an increase of the spring constant of an individual alpha helix, *κ_α_*, as shown in the Monte-Carlo simulated force-strain curves in Fig. 4a.

An increased stiffness can furthermore be explained if we regard the filament as a bundle of protofilaments. If these protofilaments are fully coupled to each other, the persistence length of the bundle increases as *N*^2^ instead of linearly in *N* in the uncoupled case. ^43^ An increase in coupling strength and thus an increased initial slope may be caused by higher salt concentration or lower pH and is thus in line with our experimental results. It should be noted, however, that we do not include this effect in our simulations. Instead, we simulate ‘subunit coupling’, see Fig. 4b, that describes the size of fully coupled lateral subunits into which the monomers are organized. ^35^ Thus, in higher order subunits, more elements have to unfold simultaneously to achieve the length change, Δ*L*, of the filament. This effect already plays a role for the initial stretching and leads to a slightly increased slope (see inset of Fig. 4b).

The origin of intra-filament coupling is found in the structure of vimentin. Each vimentin monomer has an excess of −19 *e* negative charges. A representation of the charge distribution in the vimentin monomer is shown in Fig. 5a. The excess negative charges are mostly found in coil 1B, coil 2 and the tail. As most polar and charged amino acids are not buried within the alpha helix ^44^, the excess negative charges of neighboring monomers within the filaments are likely to be in close proximity. The resulting repulsion presumably destabilizes the filament structure. The decreased initial slope and reduced stability we observe in low salt conditions therefore indicates that cations screen or – in case of multivalent ions – cross-bridge these repulsions. This is supported by the fact that vimentin is known to assemble in the presence of a sufficiently high concentration of cations and to disassemble in low salt buffer. ^36^

**Fig. 5.**
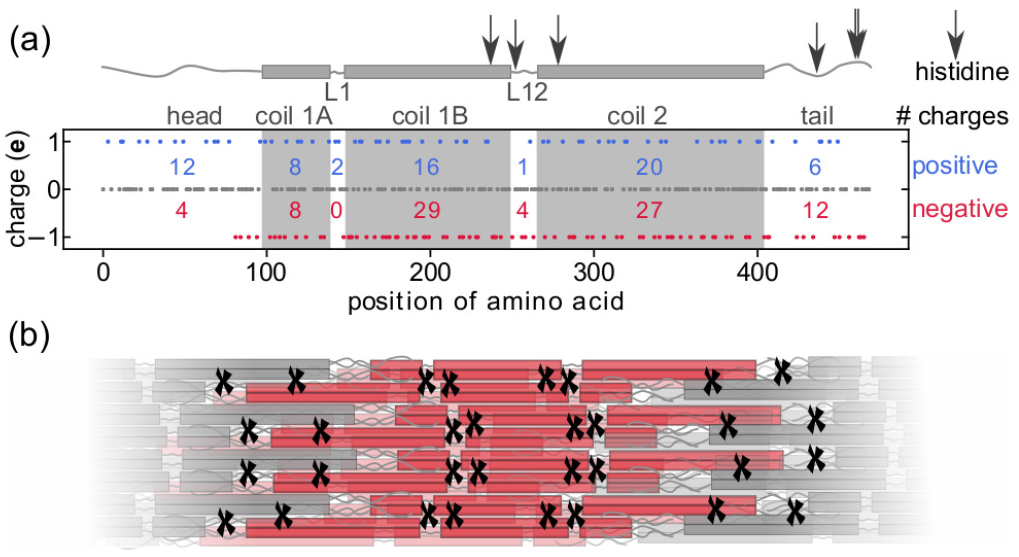
Charged amino acids in vimentin. (a) Top: Schematic representation of the vimentin monomer. The positions of histidines in one vimentin monomer are marked with arrows. Bottom: The charges of the amino acids are plotted versus their position in the sequence. The number of amino acids that carry a positive (lysine and arginine, blue) or negative charge (aspartic acid and glutamic acid, red) at pH 7.4 are shown for the head, coils, linkers and tail. (b) The possible additional intra-filament interaction sites upon charge changes in the histidines are marked by ‘*x*’ for the visible front half of the red ULF.

In contrast to the ion mediated, indirect, electrostatic effect, a pH change directly influences the charge pattern of the protein sequence. To be specific, in the pH range used here, we switch histidines from being uncharged to positively charged in an acidic environment. ^45,46^ In vimentin there are six histidines at various positions, indicated by the arrows in Fig. 5a. These additional positive charges increase the number of possible electrostatic interaction sites between monomers in the filament (Fig. 5b), and therefore in our case decrease the repulsion, increase attraction, and promote coupling in the filament, similar to added ions.

Whereas the change of the initial slope is well explained by variations of *κ_α_*, other experimentally observed changes in the force-strain curves, such as the plateau slope and length are not reproduced by this variation. To be able to compare the mechanisms that affect the plateau, we first examine the unfolding reaction that leads to the plateau formation. The force level of the plateau *F*_plateau_ (Fig. S10†), the force reached at strain *ε*_I_, is a measure of the energy necessary for unfolding. Fig. 4c shows a schematic and simplified energy landscape for the transition from the alpha to the unfolded state (green). The energy barrier *E_A_* between the two states is indicated in Fig. 4d. The optical trap is approximated by a harmonic potential (dashed line). By applying a force (Fig. 4d), the harmonic potential is moved to the right, thereby decreasing the energy barrier in the total potential (blue) and the unfolded state becomes more probable.

The simulated force-strain curves in Fig. 4b reveal a strong dependence of *F*_plateau_ on the subunit size. By choosing a small subunit size such as *N_sub_* = 4 we are able to reproduce the observed decrease of *F*_plateau_ in low salt buffer. At low pH, *F*_plateau_ is even higher than in high salt buffer. Thus, *EA* is apparently even further increased at low pH. This behavior may be explained by an increased free energy difference, Δ*G*, between the alpha and unfolded state (purple curve in Fig. 4h as compared to green curve in Fig. 4d), thus rendering the transition from the alpha to the unfolded state less probable at the same applied force (Fig. 4g,h), effectively increasing *F*_plateau_. Fig. 4e shows how Δ*G* influences the force-strain curve.

In addition to an increased *F*_plateau_, we also observe a shortening of the plateau at low pH (Fig. 2b). The length of the plateau, *ε*_II_ *−ε*_I_, depends on the number of unfolding events and the length increase during unfolding, Δ*L*. Here, by the ‘number of unfolding events’ we summarize (i) fully unfolded ULFs and (ii) partially unfolded ULFs, as each of them consists of 32 monomers with three coils each, which can unfold fully or in parts. As *ε*_I_ is relatively constant in all measuring conditions, *ε*_II_ is a measure for the length of the plateau. Fig. 4f demonstrates how a decrease of Δ*L* shortens the plateau.

Earlier interpretations of the plateau being a transition from the alpha helices to beta-sheets would allow for an elongation to strain 0.77^8,18^ in the plateau. This value agrees with *ε*_II_ at high salt concentrations, but is exceeded at low salt conditions and not reached at low pH values (Fig. 2). Recent results indicate that the unfolding is in fact not a two-state process but that alpha helices first unfold to a random coil structure. ^19^ These random coils could be either longer or shorter than the beta-sheet conformation and thereby explain the variations in Δ*L*. The remarkably short plateaus we observe at low pH indicate a strong influence of the pH on Δ*L*. From the simulation in Fig. 4f we learn that decreasing Δ*L* furthermore increases the slope of the plateau, which agrees well with the increase of the slope we observe at low pH (Fig. 3c). The additional positive charges located at the sites of the histidines might act as crosslinkers in the filament, ‘locking’ the monomers in place and thereby decreasing Δ*L*.

Combining the simulations and the experimental results, we are now able to explain the observed increase of the initial slope, shortening of the plateau, shift to higher forces *F*_plateau_ at low pH or high salt conditions by increased subunit coupling and decreased Δ*L*. Additionally, for the low pH conditions, a more pronounced Δ*G* comes into play, whereas for high salt, *κ_α_* is increased. The slope of the plateau can be modeled by a decrease of the subunit size or of Δ*L*. As we observe no change of the slope of the plateau throughout all salt conditions at pH 7.5, these effects seem to be balance out during the plateau formation.

Previous studies of the stability of single coiled coils at varying pH and ionic strength showed seemingly conflicting results. ^22,24,26^ These opposing observations can be understood in the light of different primary sequences that lead to coiled coils of varying stability, based on the length of the coils, ^21^ the hydrophobic packing or helix propensity, *i.e.* how prone an amino acid is to form alpha-helical structures, ^27^ and electrostatic interactions in the coil. ^22,24,26^ It is, however, remarkable that even for coiled-coils, which have a very defined amino acid pattern, the response to external charge cues can vary to the reported extent. The effect of the electrostatic environment we report here is valid for vimentin but might be considerably weaker or stronger for other types of IFs. As the charge patterns in different types of IFs vary, we expect the extent of the effect of salt ions to scale according to the strength of the repulsion between subunits in the different IFs. For example, similar IFs, such as desmin and glial fibrillary acidic protein (GFAP) have an excess charge of −15 *e* and −13 *e*, respectively. We therefore anticipate the effect of ions to be weaker in these filaments. For the nuclear IF lamin A, which has an excess charge of −3 *e*, ^1^ the dependence of the mechanical properties on ion concentrations is likely to negligible. For the pH dependence, we expect the impact to scale with the number of histidines per monomer rather than the excess charge of the protein sequence. Whereas vimentin and desmin posses six histidines (*N_H_* = 6), GFAP (*N_H_* = 8) contains more potential additional interaction sites at low pH. ^1^ The effect in GFAP may therefore be even stronger than observed here for vimentin. For lamin A, *N_H_* = 13, which leads us to the hypothesis that lamin A is more sensitive to pH variation than vimentin but less prone the mechanical changes by variations of salt concentration. The exact pH values the different IFs are sensitive to depend on the pKa of each single histindine.

As the cytoplasmic pH typically lies between 7.0-7.5, ^41^ the mechanical properties of vimentin are susceptible to pH changes in this range. Thus, cells are equipped with a “tool” to rapidly and locally tune their stiffness without remodeling the whole cytoskeleton. However, it remains unclear, how relevant the adaptability of the IF mechanics is in living cells. One intriguing phenomenon, where considerable pH changes and the up-regulation of vimentin in epithelial cells coincide, is wound healing. In epithelial wounds, mesenchymal cells expressing vimentin promote healing of the skin. During the restoration of the tissue, these cells might be exposed to pH milieus reaching from the healthy skin pH (4.0-6.0) to the body’s internal pH (7.0-7.4). ^42,47^ Vimentin is not only up-regulated in the cells but also excreted into the extracellular space. ^48^ The result of our studies suggest that the mechanical changes vimentin undergoes in this pH range are considerable and might play a role within cells as well as the extracellular space during skin repair. Direct studies of the mechanical properties of cells and the corresponding structure of the IF network after exposure to varying pH milieus are necessary to make a more substantiated statement about the role of pH milieus for IF mechanics. Intracellular and extracellular pH gradients have furthermore been found to be a feature of migrating tumor cells. A decrease of the pH along the axis of migration as well as local pH variations, such as nanodomains of high pH close to focal adhesions, have been reported. ^49^ An influence of the pH on the actin cytoskeleton is known, ^49^ however, the role of IFs in this scenario is so far, to our knowledge, unexplored.

IFs have been shown to provide a “rescue mechanism” for cells that are subject to large areal strains. ^40^ At high strains, the actin network cannot compensate the applied force and IFs become load bearing. The extensibility of IFs therefore seems to be a crucial filament property, allowing them support the cell at these high strains. In the case of decreased pH, IFs are extended less at equal force, potentially leading to network stiffening. According to the strong effect we observe, we speculate, that local, intracellular pH changes may influence the role the IFs play under strain considerably. Studies of the strain response of single cells and multi-cellular ensembles at varying cell internal pH may shed light onto the role of IFs for cell mechanics in general. As a change of the pH milieu is often correlated with stress, the results for single filaments presented h ere, together with future experiments in cells may improve our understanding of biological stress response.

## 4 Conclusions

To conclude, we directly relate the mechanical response of single vimentin filaments to stretching in different buffer conditions to variations in the molecular electrostatic interactions in the filament. Our results show that the strong response to the electrostatic environment reported for coiled coils is preserved in mature vimentin filaments. A likely interpretation is that salt ions in the buffer screen or bridge electrostatic repulsion in the hierarchical structure and thereby stabilize the filaments. Additional positive charges in the amino acid sequence caused by a lowered pH stabilize and stiffen vimentin filaments as well. Thus, our results indicate that the mechanical role of IFs in cells can adapt to local pH and ion concentrations. Both effects, salt and pH, may allow cells to locally tune their stiffness without having to rebuild the entire cytoskeleton and thereby adapt their mechanics to varying requirements. In this context, we show that stiffening of vimentin networks that was previously reported upon the addition of Mg^2+^ relies on increased inter-filament interactions and does not originate form stiffening of single filaments. Thus, by ensuring a relatively constant stiffness, extensibility, force-strain behavior, and stability of the filaments at physiological potassium concentrations and in conditions that are known to affect the bundling behavior of vimentin, we suggest that network mechanics can be tuned independent of the single filament properties. Consequently, the next step is to study, how the variability of single filament mechanics translates to network properties. This will allow for relating the intra-filament interactions studied here and inter-filament interaction in networks.

## Conflicts of interest

There are no conflicts to declare.

## Acknowledgements

We thank J Forsting, J Kraxner, H Herrmann and JC Thiele for helpful discussions, S Bauch and K Sabagh for technical support, U Rölleke and H Somsel for critical reading of the manuscript.

This project was funded by the European Research Council (ERC) under the European Union’s Horizon 2020 research and innovation program (Consolidator Grant Agreement no. 724932). This research was conducted within the Max Planck School Matter to Life supported by the German Federal Ministry of Education and Research (BMBF) in collaboration with the Max Planck Society. The work further received financial support via an Excellence Fellowship of the International Max Planck Research School for Physics of Biological and Complex Systems (IMPRS PBCS) and the Studienstiftung des deutschen Volkes e.V..

## Notes

### Competing Interest Statement

The authors have declared no competing interest.

### Summary of Updates

Minor changes.

