## Supplementary Information for "Tuning Intermediate Filament Mechanics by Variation of pH and Ion Charges"

### Electronic Supplementary Information (ESI) for:

### Flow simulations

The concentrations and pH conditions given are the buffers that were injected into the microfluidic chip. Because the flow was stopped during incubation and stretching of the filaments, diffusion of the cations between the assembly and measuring buffers, and the assimilation of the pH have to be considered. This means that the conditions in proximity to the filament during the measurement were slightly different to the injected buffer. The temporal evolution of the salt concentrations and the pH at the measurement position was simulated and is described in the following.

#### Methods

A simplified microfluidic chip design (Fig. S9a) was used for finite element method (FEM) simulations with COMSOL Multiphysics 5.3 (COMSOL GmbH, Göttingen, Germany). Flow and diffusion were simulated for an average velocity of 0.001 m/s laminar inflow for each inlet for water with $K^+$ ($D_{\mathrm{K}} = 1.67\ 10^{-9}\ \mathrm{m^2/s}$) [1] or $Mg^{2+}$ ($D_{\mathrm{Mg}} = 0.594\ 10^{-9}\ \mathrm{m^2/s}$) ions. [1] For the pH, the concentration of hydrogen ions $c(\mathrm{H}^+)$ was calculated as

$$c(H^+) = 10^{\text{-pH}} \tag{1}$$

and the diffusion of $H^+$ was estimated by ($D_H = 8.17\ 10^{-9}\ \mathrm{m^2/s}$). [1] First, the equilibrium ion distribution in the chip was simulated under flow. Taking this as a starting condition, a second simulation was calculated without flow, only allowing diffusion. The change of the concentrations of the cations and $H^+$ ions was simulated at the position of the force-strain measurement (Fig. S9a, red mark) for a duration of 5 min. The equilibrium pH is reached within the first minute and then stays relatively constant over time and in the flow cell. This resulting equilibrium pH was clearly distinguishable for the different measuring buffers. The change of cation concentrations ($Mg^{2+}$ from the measuring buffer, $K^+$ from the adjacent assembly buffer) is shown in Fig. S9b for the example of the 10 mM $MgCl_2$ measuring buffer.

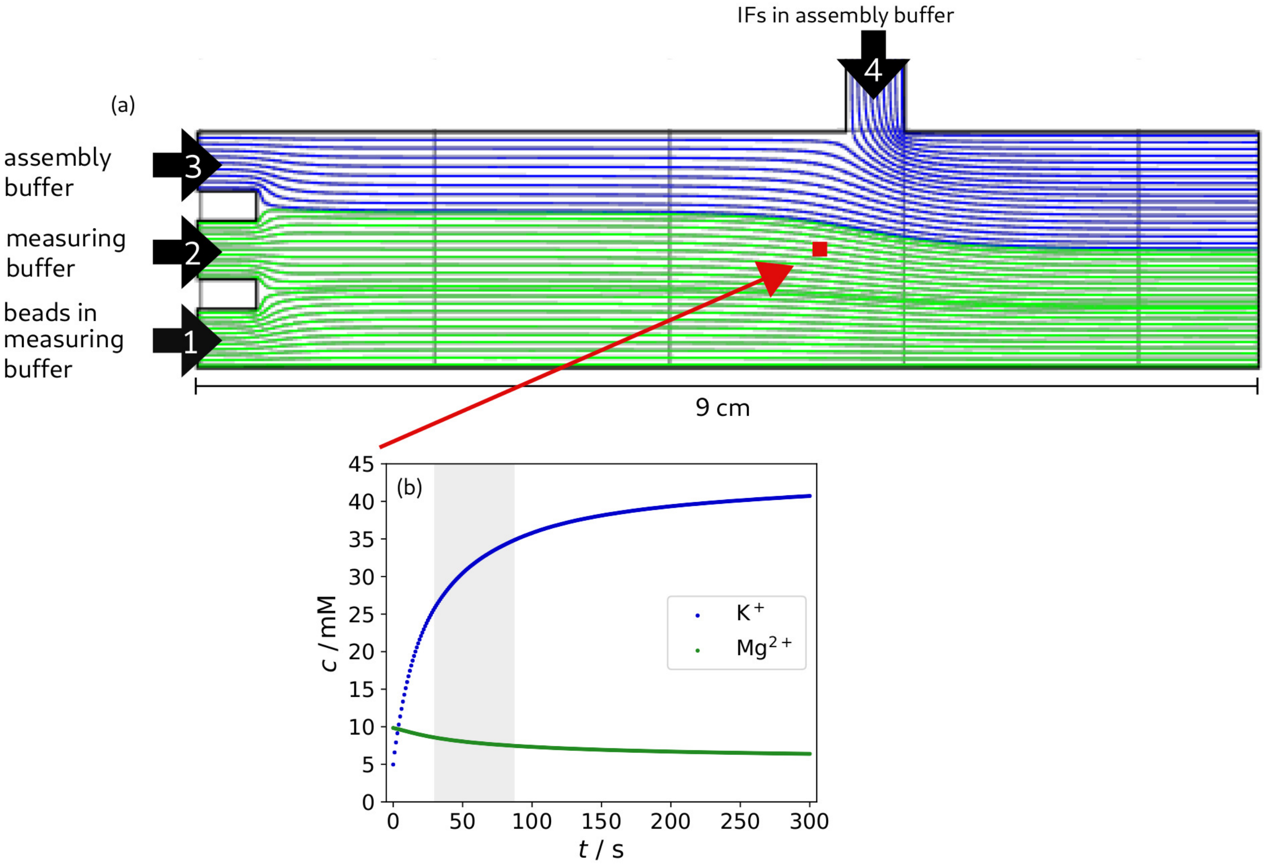


**Fig. S1 Results of FEM simulations.** (a) Schematic of the flow cell including simulated stream lines of the in-flowing buffers. In the experiment, beads in measuring buffer are injected in channel 1, measuring buffer in channel 2, assembly buffer in channel 3 and vimentin in assembly buffer in channel 4. In Fig 1b in the main text, we omit channel 2, as it is solely used for calibration of the traps and not crucial for the understanding of the measurement procedure. It is, however, relevant for the flow properties presented here. The colors correspond to the cation species of the buffer (blue: $K^+$, green: $Mg^{2+}$). For this simulation, the measuring buffer contained 10 mM $Mg^{2+}$ and the assembly buffer 100 mM $K^+$. IFs and beads were not included in the simulation. The position of the measurement is marked in red and corresponds to the position for which the development of the cation concentrations after stopping the flow was calculated. (b) Plot of the temporal evolution of the concentrations of $Mg^{2+}$ and $K^+$ ions at the measurement position after stopping the flow. The time window of the measurement is indicated.

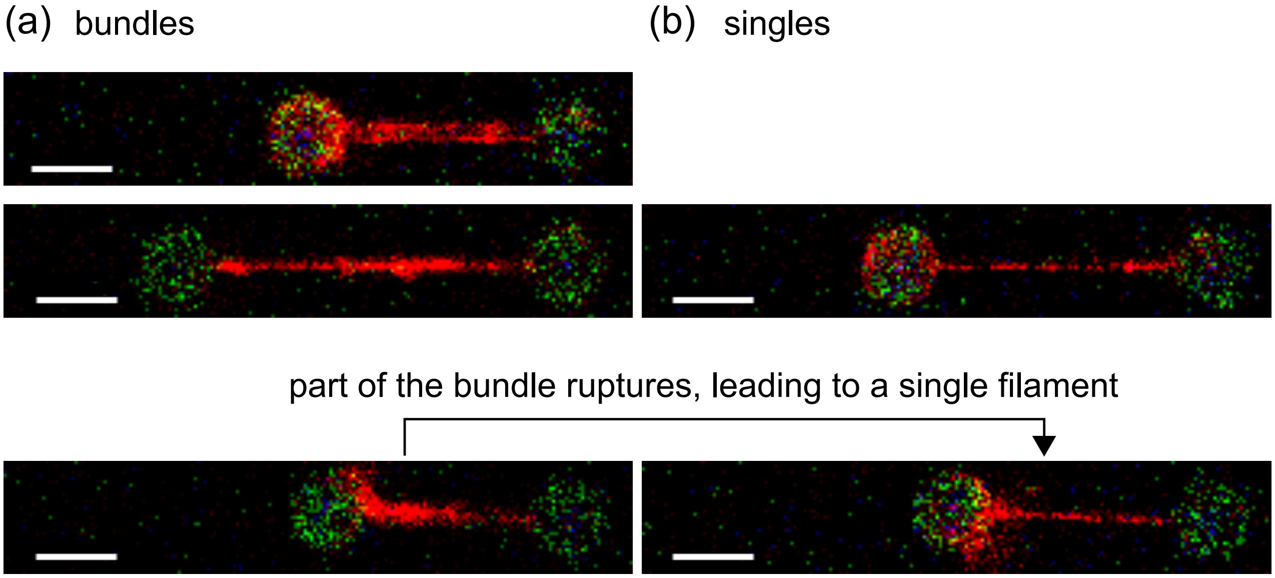


**Fig. S2 Confocal images of single filaments and bundles.** (a) Examples of vimentin IF bundles. The number of filaments in each bundle cannot unambiguously be determined. The inhomogeneity along the bundle further complicates the accurate description of the bundle. In some cases, individual filaments in the bundle rupture during the experiment, leading to one single filaments left between the beads. (b) Examples of single filaments. The scale bars are 5 μm.

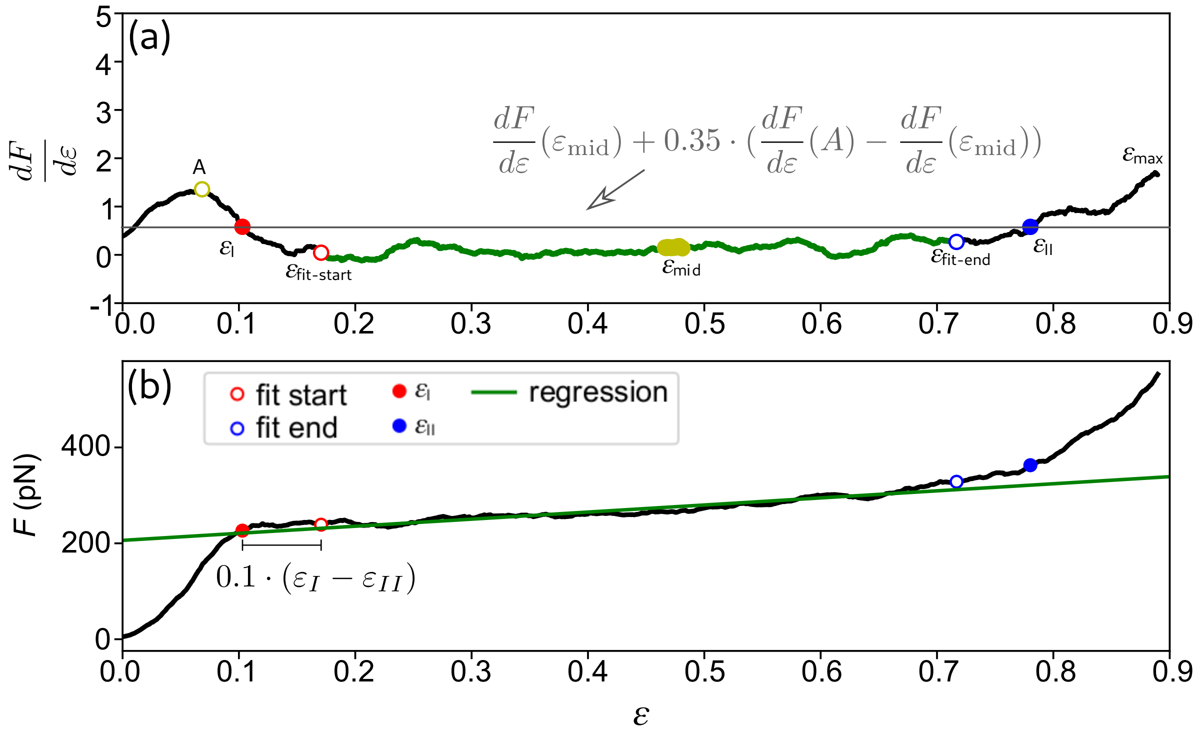


**Fig. S3 Analysis of the slope of the plateau for one single force-strain curve.** (a) Differential force-strain curve, smoothed with a moving average with the window width of $\frac{1}{20}$ of number of data points in the curve before $\varepsilon_{\mathrm{max}}$. The center of the curve ($\varepsilon_{\mathrm{mid}}$) is marked in yellow. The peak of $\frac{dF}{d\varepsilon}$(A) is shown with the open yellow circle. The threshold for the plateau is indicated with the grey line. The values for $\varepsilon_{\mathrm{I}}$ (red, solid) and $\varepsilon_{\mathrm{II}}$ (blue, solid) at the threshold are shown. The open red and blue circles flank the region used for the calculation of the regression (green). (b) Resulting regression plotted together with the raw force-strain curve.

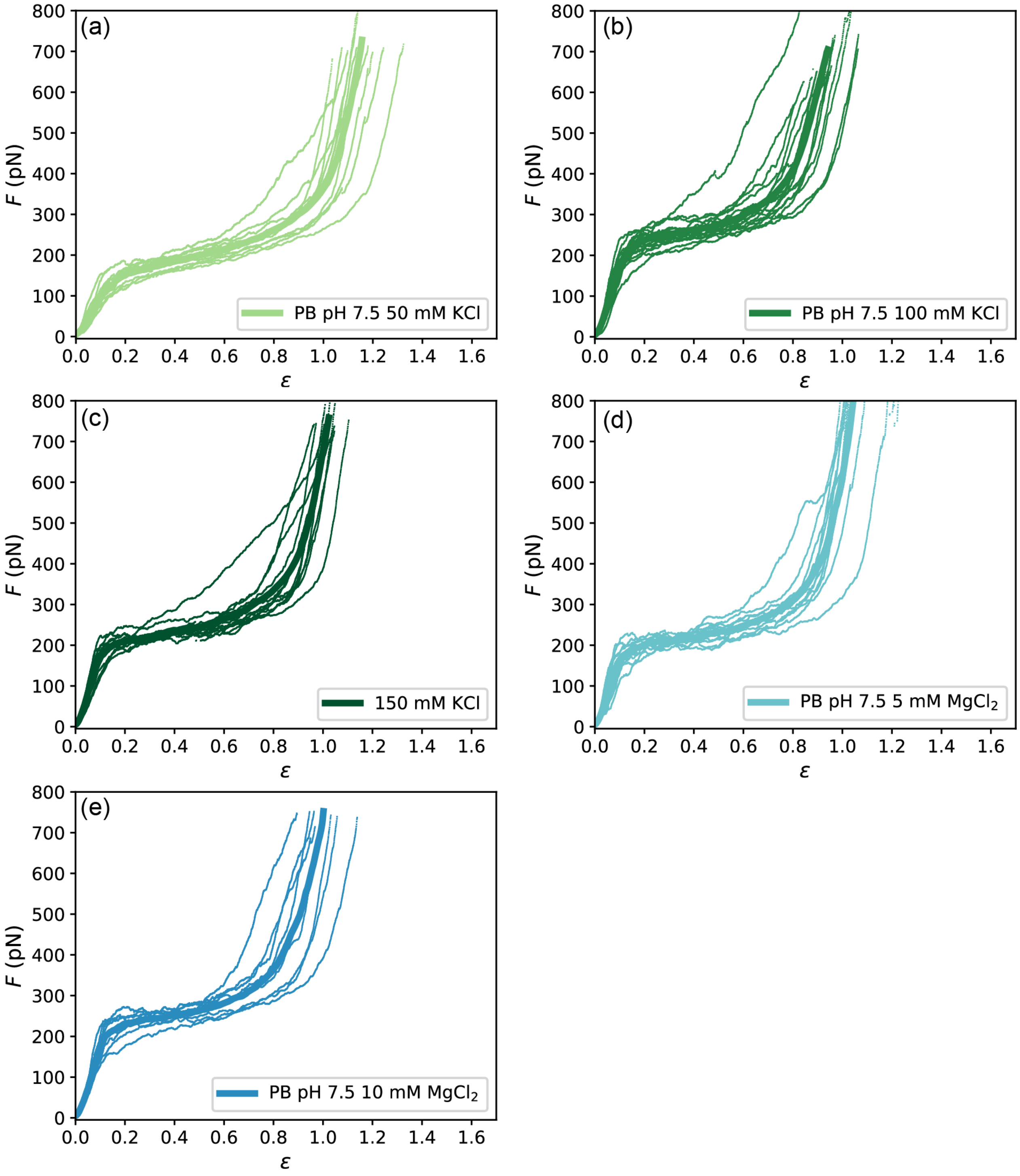


**Fig. S4 Force-strain curves for each salt condition measured.** (a)-(e) All single measurements of meta stable and stable filaments are plotted (thin lines) along with the average curves (bold lines, as shown in Fig. 2a in the main text).

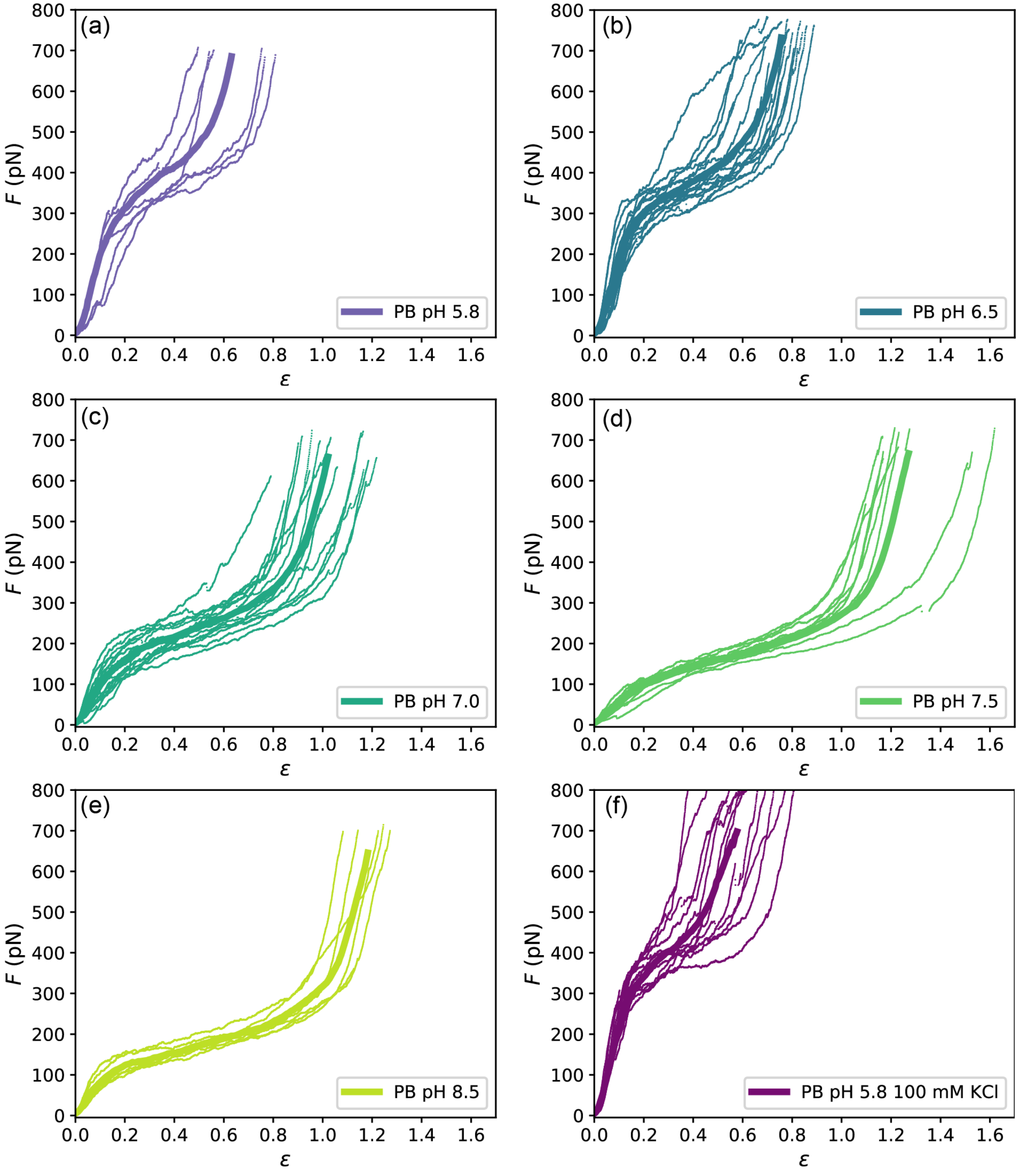


**Fig. S5 Force-strain curves for each pH condition measured.** All single measurements of meta stable and stable filaments are plotted (thin lines) along with the average curve (bold lines, as shown in Fig. 2b) (a)-(e) Show data recorded at increasing pH values and (f) measurements at pH 5.8 with 100 mM KCl.

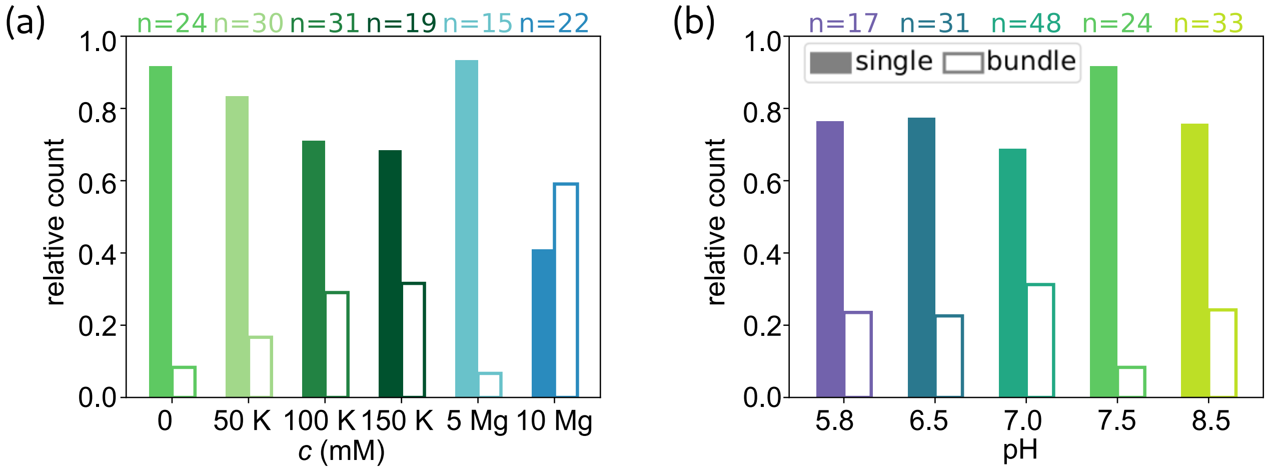


**Fig. S6 Relative count of single filaments and bundles for each condition.** The bundles were identified from the confocal images and from the force data. Only single filaments were used in further analysis. (a) The increased fraction of bundles at higher salt concentrations indicates that the ions promote filament bundling. (b) The fraction of bundles does not show a trend at different pH conditions.

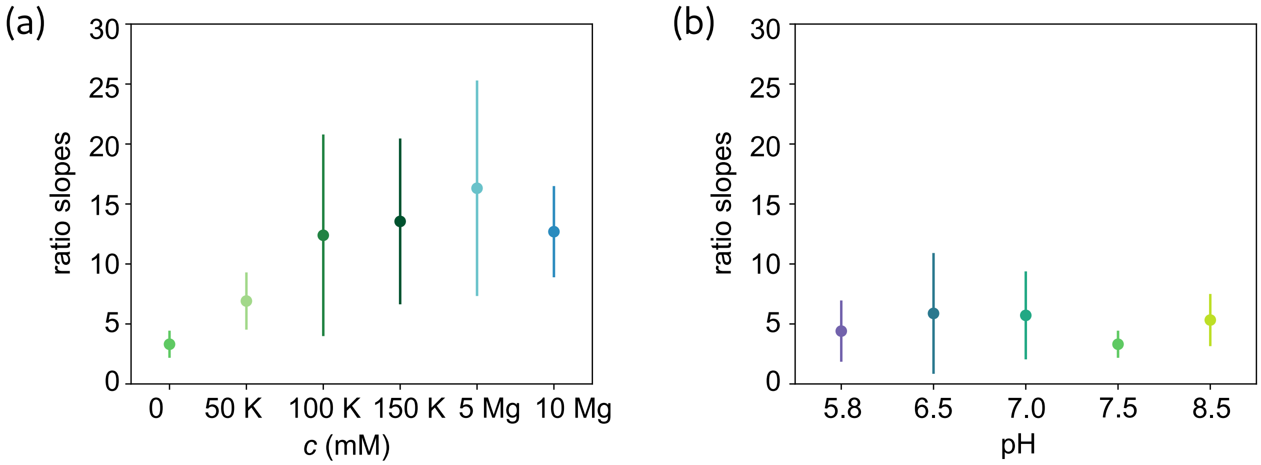


**Fig. S7 The ratio of the initial slope and the slope of the plateau for each measuring condition.** The respective slopes are presented in Fig. 3 in the main text. (a) The ratio changes at different salt conditions as the plateaus have the same slope, whereas the initial increase changes at different salt conditions. (b) The ratio of the slopes in buffers of increasing pH stays relatively constant.

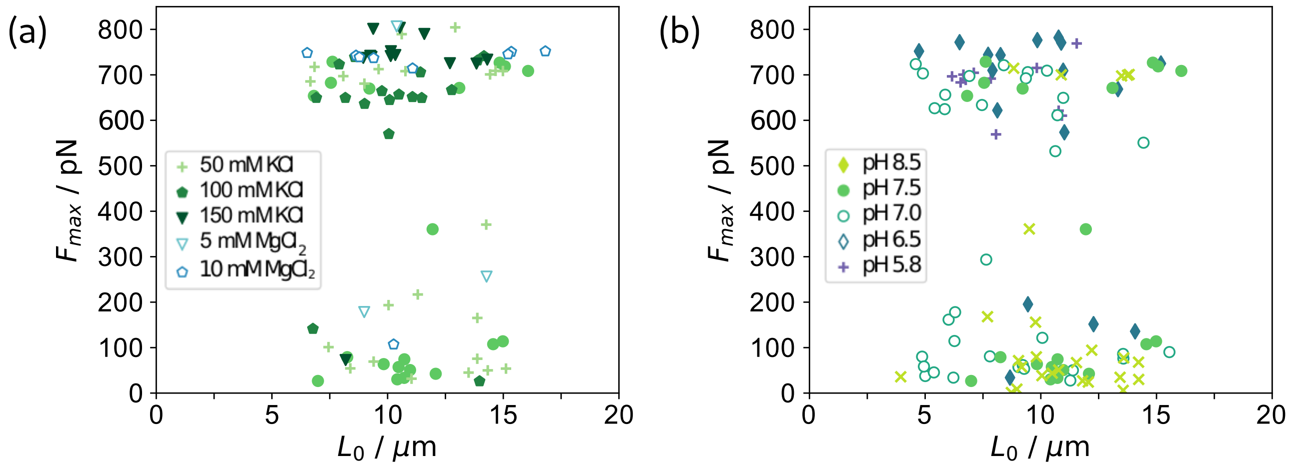


**Fig. S8 Maximum force plotted against initial filament length for each filament.** There is no correlation between the initial filament length and the maximum force reached during stretching. The data cluster at forces around 700 pN because beads are pulled out of the traps. (a) For varying salt concentrations, (b) for varying pH conditions.

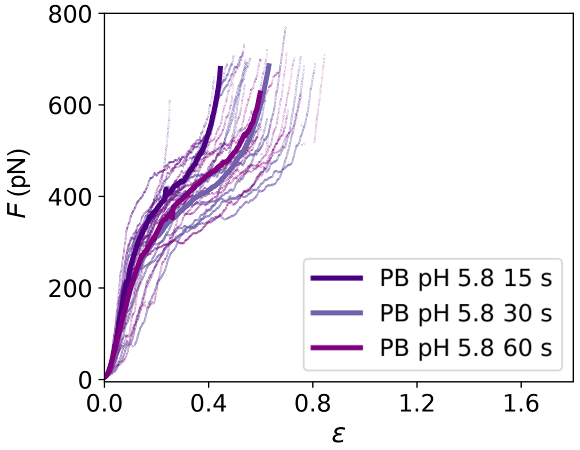


**Fig. S9 Force-strain curves for filaments stretched in phosphate buffer (PB), pH 5.8 with varying incubation times.** All single measurements are plotted by thin lines, the average curves are shown by bold lines. Within the variation of the single curves in each condition there is no difference apparent between the incubation times.

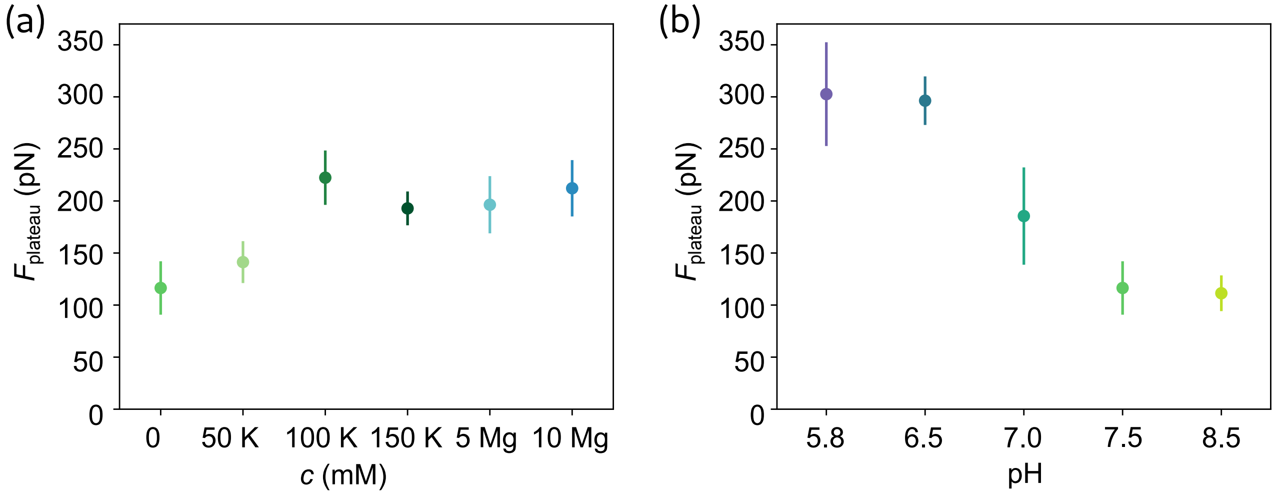


**Fig. S10 Plateau force of the force-strain curves.** The force at $\varepsilon_I$ is shown for all single force-strain curves for each condition. The error bars correspond to the standard deviation. (a) $F_{\text{plateau}}$ from curves recorded at pH 7.5 with varying salt concentrations. The plateau is shifted to higher forces for higher KCl concentrations or in the presence of $MgCl_2$. (b) $F_{\text{plateau}}$ extracted from the force-strain curves measured at varying pH values. The plateau appears at higher forces for lower pH conditions.
